# Trait – climate relationships in *Themeda triandra*, a widely distributed C4 grass and crop wild relative

**DOI:** 10.64898/2026.03.04.709158

**Authors:** Vinod Jacob, Brian J Atwell, Luke Yates, Rachael V Gallagher, Emma Sumner, Travis Britton, Ian J Wright

## Abstract

- Quantifying relationships between traits and climate using plants collected from diverse climatic origins, grown under common conditions, potentially provides valuable insights into climate adaptation.
- We report on fifteen accessions of kangaroo grass (*Themeda triandra*), a C4 species distributed across Australia, Asia, the Middle East and Africa from the Andropogoneae clade of grasses that is vital to global agriculture. Plants were grown to maturity in glasshouses under two thermal regimes, with ample water supplied. Numerous physiological, “economic” and developmental traits were characterised.
- As expected, plants grown at 20°C maxima had lower photosynthetic rates (A_sat_) and dark respiration rates, reduced leaf expansion, and delayed flowering compared with plants grown at 30°C. However, surprisingly few traits varied with climate-of-origin: accessions from colder climates had higher A_sat_ alongside lower leaf mass per area, but only when grown at 20°C; flowering time showed the strongest correlation with site climate, with plants from wetter, warmer or less variable climates taking longer to flower.
- Our findings highlight remarkable phenotypic flexibility in key traits of *T. triandra*; this flexibility is likely key to its wide distribution. The strong relationship between flowering time and climate-of-origin underscores the importance of reproductive phenology as an adaptive trait.

## Introduction

Kangaroo grass (*Themeda triandra* Forssk.) is a perennial C4 (NADP–ME) tussock grass within the Andropogoneae tribe, a speciose clade of grasses with mostly tropical and sub-tropical distribution that includes globally important crop species such as sorghum, maize, and sugarcane. *T. triandra* likely radiated from Sri Lanka to its current distribution throughout South and Southeast Asia, the Middle East and Africa (Dunning et al., 2017). *T. triandra* has also adapted to every biome throughout the Australian continent, ranging from the semi-arid interior savanna and grasslands, to the seasonally wet tropics and sub-alpine, cool temperate regions (Hayman, 1960; Gallagher, 2016; Arthan *et al*., 2022). It is therefore adapted to a wide range of temperature and rainfall regimes, degrees of seasonality, and to a variety of soil types. Its occurrence in cool climates is particularly notable because C4 photosynthetic physiology generally characterises grasses adapted to hot and dry conditions (Luo *et al*., 2024). Despite its natural range across the Australian continent and beyond (Gallagher, 2016), *T. triandra* populations have suffered dramatic declines due to land conversion for agriculture and urban development. As a result, the re-establishment of *T. triandra* populations is an important first step for the restoration of many of Australia’s grassland ecosystems (Cole & Lunt, 2005). However, *T. triandra* has limited dispersal potential hence its response to climate change is largely dependent on trait plasticity and standing genetic variation for adaptation *in situ* (Ahrens et al 2020).

Adaptation of kangaroo grass across its vast environmental range is seemingly underpinned by substantial within-species genetic variation and genome duplication (Snyman *et al*., 2013; Godfree *et al*., 2017; Ahrens *et al*., 2020; Dunning *et al*., 2022). The species comprises a polyploid complex, with diploids generally more common in the cooler, wetter areas of southeast Australia and tetraploids more common in the warm, arid areas of central and western Australia (Hayman, 1960; Ahrens et al., 2020). Previous studies on *T. triandra* that focussed primarily on population genetics (Hayman, 1960; Dunning et al., 2017, 2022) showed that polyploidy reduces the negative impacts of drought and heat stress on plant fitness (Godfree et al., 2017; Ahrens et al., 2020). However, little is known about the mechanisms that underpin the success of *T. triandra* across its vast geographic range. For example, it is unknown whether populations from contrasting climates differ predictably in key photosynthetic or hydraulic traits (or temperature tolerances thereof) and, if so, whether the trait differences are fixed (expressed similarly irrespective of growth conditions) or exhibit a high degree of phenotypic flexibility. These questions are central to understanding the success of this species throughout Australia. In principle, identifying the adaptive solutions that *T. triandra* employs to survive and thrive will not only provide new, fundamental biological understanding of this important species but also knowledge of physiological mechanisms that may be key to breeding crops that are more resilient to climate change, given its close relatedness to important crops in the Andropogoneae,. Furthermore, these insights could help guide the selection of locally-adapted genotypes to enhance the climate readiness of restorative plantings.

Climate is a major driver of species adaptation, often resulting in plant populations with distinctive phenotypes (Lowry *et al*., 2008; Lovell *et al*., 2021; James *et al*., 2023). Systematic examination of plants selected from diverse climates and grown in common environmental conditions provides insights into adaptation and the role of climate in driving coordination among functional traits (Geber & Griffen, 2003; Huxman *et al*., 2004; Albert *et al*., 2011; Schwinning *et al*., 2022). Such “common garden” experiments have revealed a variety of trait-climate patterns, across a range of species. For instance, within-species variation in net photosynthesis (A_sat_), stomatal conductance (g_s_) and leaf nitrogen (N) varied with temperature and precipitation at climate-of-origin (COO) such that plants adapted to lower temperatures had higher rates of photosynthesis and respiration when grown under common conditions (Benowicz *et al*., 2000; Ishikawa *et al*., 2007; Atkin *et al*., 2015; Zaka *et al*., 2016; Sandel & Low, 2019; Kelly *et al*., 2020). By contrast, other studies have reported no such relationship (Marchin *et al*., 2008; Aspinwall *et al*., 2013). Growth rate may be positively related to mean annual temperature (MAT) at origin (Oleksyn *et al*., 1998; Aspinwall *et al*., 2013) and negatively corelated with mean annual precipitation (MAP) (Marchin *et al*., 2008; Sandel & Low, 2019). Both a positive (May *et al*., 2017) and negative (Aspinwall *et al*., 2013) relationship has been found between flowering time and MAT, but this relationship is confounded by the impact of photoperiod on flowering (Quinby, 1972; van Esbroeck *et al*., 2003). Leaf mass per unit area (LMA) has been shown to increase with temperature (Aspinwall *et al*., 2013; May *et al*., 2017) and decrease with precipitation (Marchin *et al*., 2008; Souza *et al*., 2018), possibly attributable to the biomechanical advantages of thick, tough leaves in dry and physically demanding situations (Wright & Westoby, 2002). Leaf nitrogen per area (N_area_) showed a negative relationship with temperature and precipitation in populations of *Fraxinus americana* (Marchin et al., 2008) potentially enhancing metabolic activity and growth rates during temperature and water limitations (Chapin *et al*., 1993; Wright *et al*., 2003; Reich *et al*., 2003).

In this study we set out to better understand traitlclimate relationships in *T. triandra* by growing a selection of 15 accessions from diverse climates-of-origin under common conditions and quantifying variation in key leaf morphological and physiological traits, and the time to flowering. All 15 accessions were grown under two thermal regimes (day-time temperatures of 20°C and 30°C ), chosen to broadly represent the cool and warm extremes of the >15°C range in MAT experienced by this species across its range in Australia (6 – 29°C). This design aimed to provide insight into the robustness of observed trait–COO relationships and the degree of trait plasticity in relation to growth temperature, helping tease apart the roles of genes (G) and environment (E), and their interaction (G x E) on trait behaviour. Home climates (COO) were quantified using four climate variables: mean annual temperature (MAT), annual summed precipitation (MAP), annual temperature range (ATR) and precipitation seasonality (PS; the coefficient of variation among monthly means). Annual means/sums (MAT, MAP) were used because (1) they are undeniably important general indicators of site climate (Woodward, 1987; Moles *et al*., 2014); (2) means also capture features of low and high extremes; (3) the chief alternative, “growing season”, is impossible to define consistently for an evergreen species with broad climate range. Range and seasonality metrics were included as temporal variation in climate affects many biological processes, e.g. flowering, phenology, resource availability, productivity and functional traits (Menzel *et al*., 2006; Cullen *et al*., 2009; Heisler-White *et al*., 2009; Pérez-Camacho *et al*., 2012; Power *et al*., 2016; Lasky *et al*., 2016; Williams *et al*., 2017; Silva *et al*., 2021; Wang *et al*., 2022; Geissler *et al*., 2023).

We tested the following hypotheses based on previous major trait studies:

1. Plants from colder sites would exhibit higher rates of photosynthesis and respiration when grown under common environments (Ishikawa *et al*., 2007; Aspinwall *et al*., 2013; Atkin *et al*., 2015).
2. Plants from drier and cooler sites would exhibit higher N_area_, reflecting adaptations that enhance the efficiency of water use during photosynthesis (at least for C3 species) (Wright *et al*., 2003; Prentice *et al*., 2014; Dong *et al*., 2017) and enable continued metabolic activity and growth under low temperatures (Reich & Oleksyn, 2004; Prentice et al., 2014).
3. Plants from drier sites would have higher LMA, as required for a long leaf lifespans in physically challenging sites (Wright & Westoby, 2002), and indirectly contributing to the expected higher N_area_. Higher LMA was also predicted in plants from colder sites, as observed globally, particularly among evergreen woody species (Wright *et al*., 2005; Wang *et al*., 2023).
4. Leaves would be narrower in plants from drier and hotter sites. Smaller leaves, having thinner and therefore less insulating boundary layers, couple more tightly with air temperature, lowering the risk of overheating when transpiration is limited under hot, dry conditions (Gates, 1965; Leigh *et al*., 2017; Wright *et al*., 2017; Baird *et al*., 2021).
5. Plants originating from colder sites would flower earlier, assuming that heat sum (growing degree days) is a key trigger for flowering and that flowering is triggered by a lower heat sum at colder sites (Quinby, 1972; van Esbroeck *et al*., 2003; Aspinwall *et al*., 2016). By the same logic, we expected plants grown at the higher temperature to flower sooner than plants at the lower growth temperature.
6. Plants from climates with higher temperature and precipitation seasonality would exhibit traits aligned with the “faster” end of the leaf economic spectrum (Wright et al 2004); e.g. lower LMA leaves with higher photosynthetic and respiration rates, allowing for greater responsiveness to temporal resource heterogeneity (Reich *et al*., 2003; Poorter *et al*., 2009).

## Methods

### Plant material

Fifteen accessions of *T. triandra* (Fig. 1A) were grown from seed sourced from suppliers including the Australian Pastures Genebank and Native Seeds Pty Ltd and Nindethana Seed Service Pty Ltd. Accessions were chosen to represent a broad range of climates of origin (Table S1; MAT ranging from 11.8 - 26.9°C and MAP ranging from 281 - 1129 mm) but were also guided by practical considerations, as only accessions with sufficient germination for adequate replication were included from an original collection of more than 30 accessions. Plants were propagated from seed in 36-mm peat pellets (Jiffy Products, Shippagan, New Brunswick, Canada) in a climate-controlled growth cabinet set at 35/20°C in an equal day/night regime, with a peak irradiance of 1000 μmol photons m^-2^ s^-1^ midway through the photoperiod. Germinated seedlings in the peat pellets were moved into 4.45-L square round plastic pots containing a 9:1 mixture of commercial potting mix (<30% fine sand, >70% pine bark; Australian Growing Solutions) and field soil (40:30:30 sandy clay loam), and placed in climate-controlled glasshouses with 14-h light at a 30/26°C day/night temperature regime. Natural light was supplemented with blue/red LED lighting, adding 600 μmol photons m^-2^ s^-1^, when the light levels in the glasshouses fell below 700 μmol photons m^-2^ s^-1^. All plants were fertilised weekly with a half-strength commercial water-soluble fertiliser (AQUASOL, Yates, Australia, 23:3.95:14 nitrogen: phosphorus: potassium, with added trace elements) for the first month of growth. Once plants were ∼1 month old, they were transferred to glasshouses for temperature treatments, with half the replicates for each accession randomly chosen and exposed to a growth temperature (T_g_) of either 30/26°C (HT) or a 20/16°C (LT) day/night temperatures. One final addition of liquid fertiliser was conducted, followed by a slow-release native-plant fertiliser (10 g Osmocote Native slow release) a week later.

**Figure 1.**
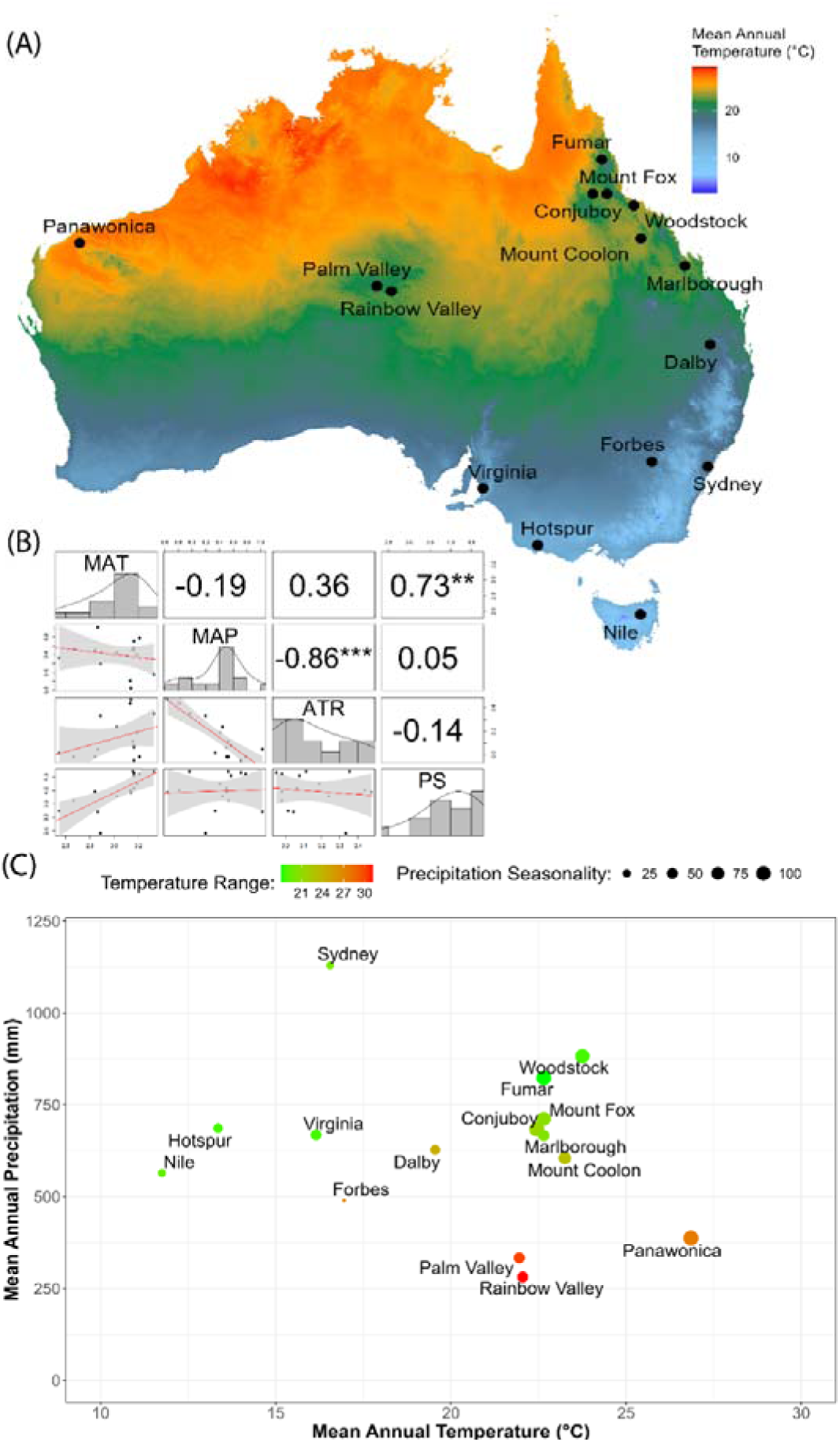
Climate-of-origin of accessions used in this study. (A) Mean Annual Temperature (MAT; °C) across Australia; accessions represented by black dots (•). (B) Pairwise correlation plots of climate predictors used in the experiment: MAT, Mean Annual Precipitation (MAP; mm), Annual Temperature Range (ATR; °C) and Precipitation Seasonality (PS, CoV). Bottom left panels show linear regressions with 95% confidence intervals. Top right panels show Pearson correlation coefficients (r) and corresponding p-values indicated by asterisks (* = p < 0.05, ** = p < 0.01 and *** = p < 0.001). Histograms along the diagonal show the distribution of each variable. (C) Scatter plot illustrating the four climatic predictors, with MAT on the x-axis and MAP on the y-axis. Dot size represents PS, while colours denote the ATR.

### Trait measurements

Tiller number, leaf length and leaf width were measured on all plants after growing for two months in the two temperature treatments. Leaf gas exchange traits, as well as leaf nutrients and LMA, were measured after two and three months growth in the HT and LT treatments, respectively, because LT plants matured more slowly than plants in the HT treatment.

Steady-state measurements of net photosynthesis (A_sat_; μmol m^−2^ s^−1^) and stomatal conductance (g_s_; mol m^−2^ s^−1^) were conducted between 09:00 and 12:00 on young fully expanded leaves using a portable open, infrared gas exchange system (LI-6800; Li-Cor, Lincoln, NE, USA) using the 1 × 3 cm aperture inserts. Measurements were conducted at treatment-specific midday air temperatures, i.e. at 20°C and 30°C for the LT and HT treatments respectively. A saturating PPFD of 1500 μmol m^−2^ s^−1^ and a chamber sample [CO ] (Ca) of 420 μmol CO_2_ mol^−1^ was maintained for both growth temperatures. Leaf-to-air vapor pressure deficit levels were maintained between 1.0 and 2.0 kPa and flow rate at 600 μmol s^-1^. Leaf area for all measurements was kept constant at 3 cm^2^ by filling the cuvette during each measurement, without overlapping of leaves. Each set of leaves was allowed ∼5 min to equilibrate before measurements were taken (Ghannoum et al., 2010). Measurements were conducted twice for each individual plant and the average values were used in subsequent analyses. Leaf dark respiration (R_dark_) was measured on the same individuals, also using the LI-6800, on the night following the measurements of A_sat_. These measurements were conducted between 23:00 and 02:00 on young fully expanded leaves, at respective night-time air temperatures (16 or 26°C for LT and HT, respectively), using a photosynthetic photon flux density (PPFD) of 0 μmol m^−2^ s^−1^, a sample [CO_2_] of 420 μmol mol^-1^, humidity maintained at ambient glasshouse conditions, and a flow rate of 500 μmol s^-1^.

Leaves analysed for gas exchange, along with others of similar age and position, were collected and scanned for leaf area (m^2^) using a Canon CanoScan LiDE300 scanner and dried at 70°C for 48 h. Leaf dry mass (g) was measured and used to calculate leaf mass per area (LMA = dry mass/leaf area; g m^-2^). Dried leaves were ground to a fine powder using a mixer mill (Retsch® MM200; Hann, Germany) and the ground sample analysed for C and N concentrations using an Elemental Analyzer (LECO TruSpec 836 Series, LECO Corporation, St Joseph, MI, USA). Leaf N content per area (N_area_) was calculated by multiplying LMA by leaf N concentration. Leaf dark respiration per mass (R_mass_) was calculated by multiplying R_dark_ by LMA. Intrinsic water use efficiency (WUE_i_) was calculated as the ratio of A_sat_ to g_s_. Photosynthetic nitrogen use efficiency (PNUE) was calculated as the ratio of A_sat_ to leaf N_area_.

### Climate-of-origin

Climate is inherently multidimensional. Rather than using a “data mining” approach as favoured in some analyses of trait-environment relationships at global scale (e.g. Joswig *et al*., 2022 used 21 climate and 107 soil variables in their analyses), here we chose to focus on four cardinal descriptors of site climate from locations (provenances) where seeds were sourced (n = 15): mean annual temperature (MAT, BioClim1m), summed annual precipitation (MAP, BioClim12), annual range of air temperature (ATR, BioClim7), and precipitation seasonality (PS, BioClim15). We used range/seasonality variables to explore the effect of variation in temperature and precipitation on plant performance. Climate data were extracted from gridded datasets from CHELSA (Karger *et al*., 2017) at a resolution of ∼ 1km at the equator and represent the long-term interpolated average of locations between 1981 - 2010.

### Statistical Analysis

We used Bayesian linear mixed models to quantify relationships between climate-of-origin (COO) and traits. All traits were log-transformed to stabilise the variances, linearise the relationships, and facilitate interpretation of the estimated effects in terms of multiplicative or relative changes. For each trait, the fixed effects comprised an overall intercept, a treatment effect, and linear terms for each of the four environmental covariates and their interactions with the treatment. To incorporate the repeated sampling structure of the data, the site of origin for each accession was specified as a random effect. To account for non-constant variance, the model for the variance included an overall intercept, a fixed treatment effect and random site effects. The prior distributions for the intercepts and variance terms were weakly informative, and the remaining fixed effects had zero-centred Gaussian priors with a common estimated variance. Use of the latter guards against overfitting, given the relatively low number of sites, and addresses any remaining multicollinearity between predictors. All models were fit in the R programming language (R Core Team, 2024) using the package brms (Bürkner, 2017). We ran four chains of 2000 iterations, including 1000 warmup iterations, and established model convergence using the improved R^"^ statistic (Vehtari *et al*., 2021). We checked the validity of the fitted models using residual plots and posterior predictive checks. In addition to plotting the posterior density for each parameter, statistical inferences for each parameter were made using the two-sided posterior probability (P_mcmc_) which is twice the proportion of the posterior samples below zero (or above zero, whichever is smallest) and closely related to the frequentist p-value associated with the two-sided null hypothesis of zero effect (Shi & Yin, 2021).

### Residualisation of seasonality predictors

We originally intended to include the climate means (MAT and MAP) in the same model as climate ranges and seasonality (annual temperature range and precipitation seasonality) but this posed serious issues for statistical inference due to the strong collinearities between them (Dormann *et al*., 2013). In this study MAT and precipitation seasonality were strongly collinear, as were MAP and annual temperature range. Rather than simply omitting annual temperature range and precipitation seasonality, we opted to residualise them with respect to MAT and MAP. Residualisation (García *et al*., 2020) means that instead of using the observed values of annual temperature range and precipitation seasonality as predictors on the model, we use their residuals from an initial linear regression of each one against MAT and MAP. This process is effective in removing collinearity, but the estimated effects for residualised variables require more effort to interpret. For example, a positive residualised value of annual temperature range at a given site means that the site has a greater range in annual temperature than would otherwise be expected given the MAT and MAP at this site. If a negative effect size is estimated for the residualised annual temperature range, this suggests that positive residualised annual temperature range values (higher than expected annual temperature range for a given MAT-MAP) are associated with a reduction in the trait value. By residualizing annual temperature range and precipitation seasonality against MAT and MAP, the latter are given the first opportunity to explain trait variation which makes their estimated effects robust to the inclusion or exclusion of the residualised predictors (Graham, 2003). The preferential treatment of MAT and MAP accords with our prior hypotheses that these are the predominant drivers of trait variation but still allows us to make useful inferences about the effect of climate ranges and seasonality which would otherwise be missed if these were dropped from the model altogether as is commonly done.

## Results

### Differential treatment effects of plants growing at 20°C versus 30°C

Averaged across all accessions, growing plants at T_g_ = 20°C (low temperature, LT) rather than T_g_ = 30°C (high temperature, HT) had a significant impact on most measured traits. Growth at LT led to a 32% reduction in A_sat_, accompanied by a 10% lower g_s_, together driving a 19% reduction in their ratio, WUE_i_ (Fig 2). Mean N_area_ (and N_mass_) did not change significantly with growing temperature, meaning that PNUE (the ratio of A_sat_ to N_area_) decreased by 35% in parallel with the decrease in A_sat_. On average, R_mass_ was reduced by 37% at the lower growth temperature. LMA and LDMC increased by 12% and 17% respectively; leaf width decreased by 11% at LT compared with HT. Leaf length and tiller number were the traits most strongly affected, decreasing by 57% and 54% respectively in cool conditions. Time to flowering was positively affected by growth temperature, taking on average 31% longer at 20°C than at 30°C (Fig. 2).

**Figure 2:**
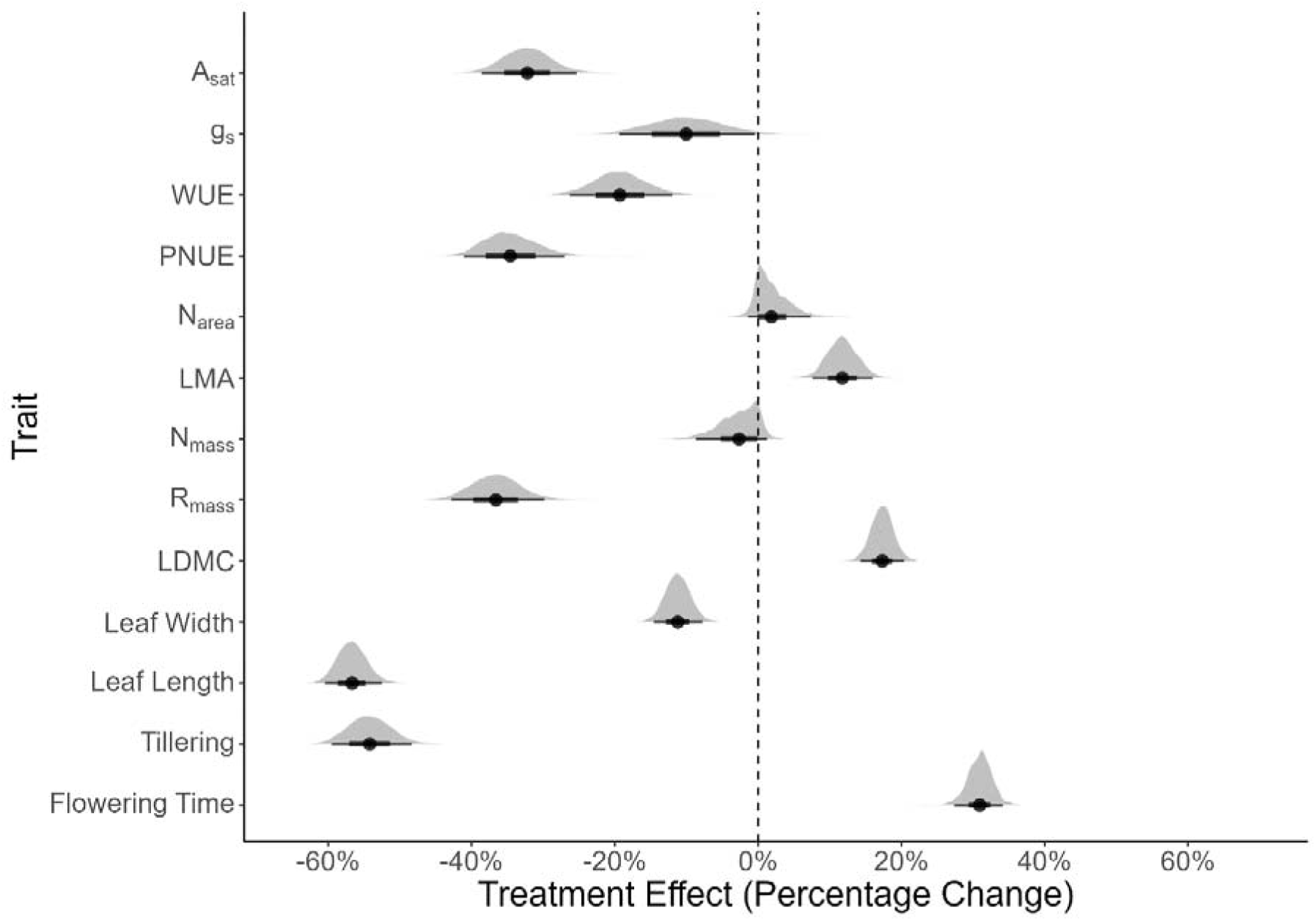
Mean effect of growth temperature treatment (hot vs cold) by trait. Abbreviations: A_sat_ = light saturate photosynthetic rate, gs = stomatal conductance to water vapour, WUE = intrinsic water use efficiency (A_sat_:g_s_), PNUE = photosynthetic nitrogen use efficiency (A_sat_:N_area_), N_area_ = lean Nitrogen per area, LMA = Leaf mass per area, N_mass_ = leaf nitrogen concentration (%), R_mass_ = leaf dark respiration per mass and LDMC = Leaf dry matter content. The x-axis denotes percentage change in the modelled trait value due to cold growth temperature (20°C) relative to the reference value of the hot growth temperature (30°C). The points are the posterior means, the thick and thin lines are the 50% and 95% credible intervals, and the grey areas are the posterior densities.

### Relationships between site climate (temperature and precipitation) and plant traits

Surprisingly few relationships were observed between site climate-of-origin (COO) and traits of *T. triandra* measured under glasshouse conditions. In general, traitlCOO relationships were generally more evident when plants were growing at 20°C (LT) than at 30°C (HT). For Figures 3-5 it is key to note that these are partial residual plots, meaning (i) they illustrate the effects of MAT with that of MAP held constant, and vice versa; and (ii) the effects of climate seasonality (ATR, PS) were quantified via residualisation, that is, as secondary effects after the influence of both MAP and MAT were accounted for.

**Figure 3:**
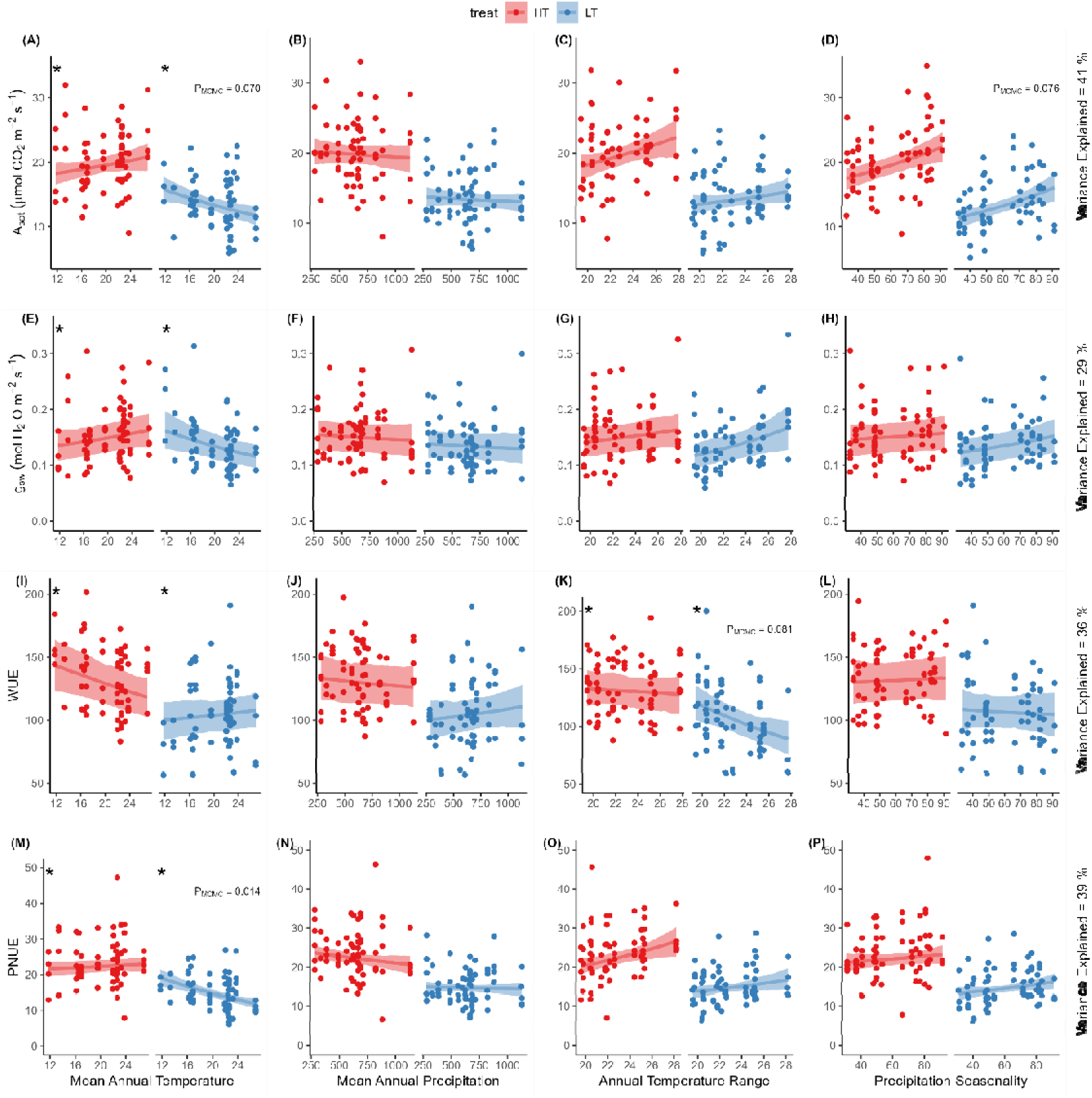
Conditional effects of site climate-at-origin on key leaf gas exchange traits. Traits: Light saturated photosynthesis (A_sat_), stomatal conductance (g_s_), intrinsic water use efficiency (A_sat_:g_s_) and photosynthetic nitrogen use efficiency (A_sat_:N_area_). The solid lines and the shaded regions denote the mean and one standard error, respectively, of the posterior predictive distribution of the fitted models. The predictive trend for each variable and treatment is conditioned on the remaining variables being held at their mean observed value. The coloured dots are the partial residuals which are the ordinary residuals added to the conditional mean. The two-sided posterior probabilities for each slope (p_MCMC_) and for each slope difference (p_Δ,MCMC_) have been indicated if they were less than 0.1. Numerical p values are displayed for slopes associated with each growth temperature and an (*) for slope differences. For all features, red and blue denote the hot and cold growth temperatures (30°C and 20°C), respectively. Detailed information on the posterior distributions of model parameters is provided in Figure S1.

Considering leaf gas exchange traits, A_sat_ (Fig. 3A) was on average higher in plants from colder sites (lower MAT), as predicted (Hypothesis 1), but only when they were grown at LT. There was no effect of MAT on A_sat_ when plants were grown at the higher temperature and, indeed, the fitted slopes at LT and HT were found to differ (wherein * in Fig. 3A denotes a p_MCMC_ value < 0.1 for the difference in slopes between the two growth temperatures). By contrast, A_sat_ was unrelated to site precipitation (Fig. 3B). Mirroring the results for A_sat_, PNUE (i.e., the ratio of A_sat_ to N_area_; Fig. 3M) was on average higher in plants from colder sites (lower MAT), but only when they were grown at LT; at HT there was no relationship. PNUE, like A_sat_, was also unrelated to MAP (Fig. 3N). Stomatal conductance to water, g_s_, was unrelated to either MAT or MAP, and at either growing temperature (Figs 3E,F). This was also the case for the ratio of A_sat_ to g_s_, WUE (Figs 3I,J).

Considering leaf economic traits, N_area_ was unrelated to site climate (Figs A,B). This was unexpected; we had predicted N_area_ to be negatively related to both MAT and MAP (Hypothesis 2). N_mass_ was unrelated to site climate (Figs 4I,J), and the same was true for leaf dark respiration rate, R_mass_ (Figs 4 M,N). This contrasts with our prediction that plants from colder sites would show higher R_mass_ when grown under common conditions (Hypothesis 1). LMA showed stronger patterning to site temperature, being on average lower in plants from colder sites when grown at LT (Fig. 4E) although unrelated to MAT for plants grown at the higher temperature (further, the LT and HT slopes were deemed to differ). LMA was unrelated to MAP at either growing temperature. Note, our hypothesis was for LMA to be higher at both drier and colder sites; i.e. it was not supported. LDMC (Fig. 4Q) showed the same pattern as LMA, being lower in plants from colder sites but only at LT, and unrelated to site rainfall.

**Figure 4:**
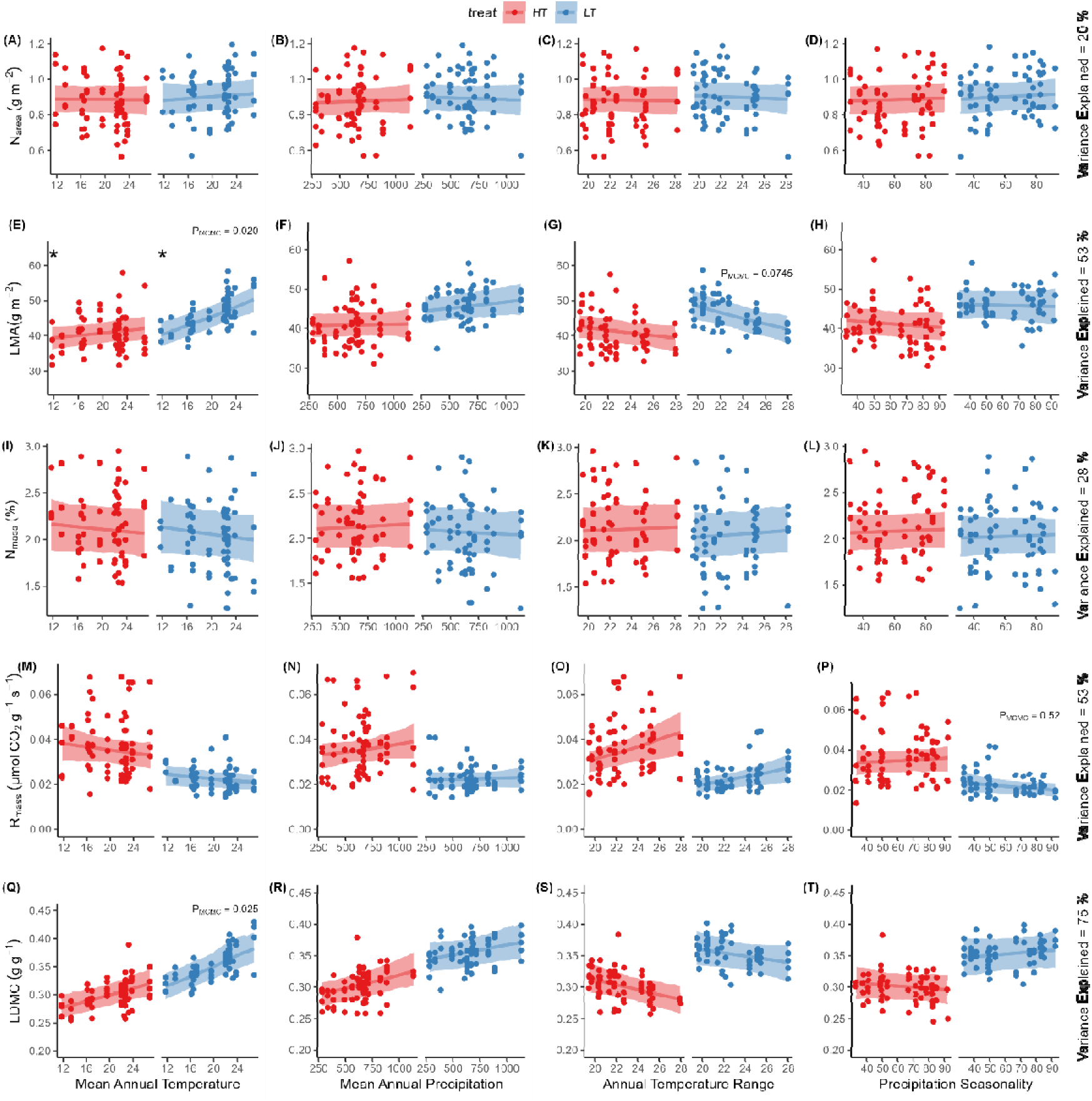
Conditional effects of site climate-at-origin on key leaf economic traits. Traits: leaf nitrogen per area (N_area_) leaf mass per area, leaf nitrogen concentration (N_mass_, %), leaf dark respiration rate per mass (R_mass_) and leaf dry matter content (LDMC). The solid lines and the shaded regions denote the mean and one standard error, respectively, of the posterior predictive distribution of the fitted models. The two-sided posterior probabilities for each slope (p_MCMC_) and for each slope difference (p_Δ,MCMC_) have been indicated if they were less than 0.1. Numerical p values are displayed for slopes associated with each growth temperature and an (*) for slope differences. For all features, red and blue denote the hot and cold growth temperatures (30°C and 20°C), respectively. For an explanation of conditional effects, refer to the caption in Figure 3. Detailed information on the posterior distributions of model parameters is provided in Figure S2.

Leaf width was unrelated to MAT at either growing temperature (Fig. 5A; in contrast to Hypothesis 4), although the LT and HT slopes were in fact deemed to differ. Also against expectations, leaf width was unrelated to MAP (Fig. 5B; we had predicted a positive relationship). Trends in leaf length differed markedly from those in leaf width in that the longest leaves were found in plants from hotter sites, at least when grown at HT (Fig. 5E); no relationship was observed at LT. Leaf length was unrelated to MAP at either growing temperature. Tiller number showed no relationship to MAT or MAP (Figs 5I,J) although in both cases the LT and HT slopes were deemed to differ. Flowering time showed the strongest climate signal of all traits. Plants from warmer environments took significantly longer to flower at both growth temperatures (as predicted: Hypothesis 5), but the relationship slope was steeper at LT (Fig. 5M), indicating a stronger temperature sensitivity for flowering at cooler growth conditions. Flowering time was longer in plants from wetter sites, but only when grown at LT (Fig. 5N).

**Figure 5:**
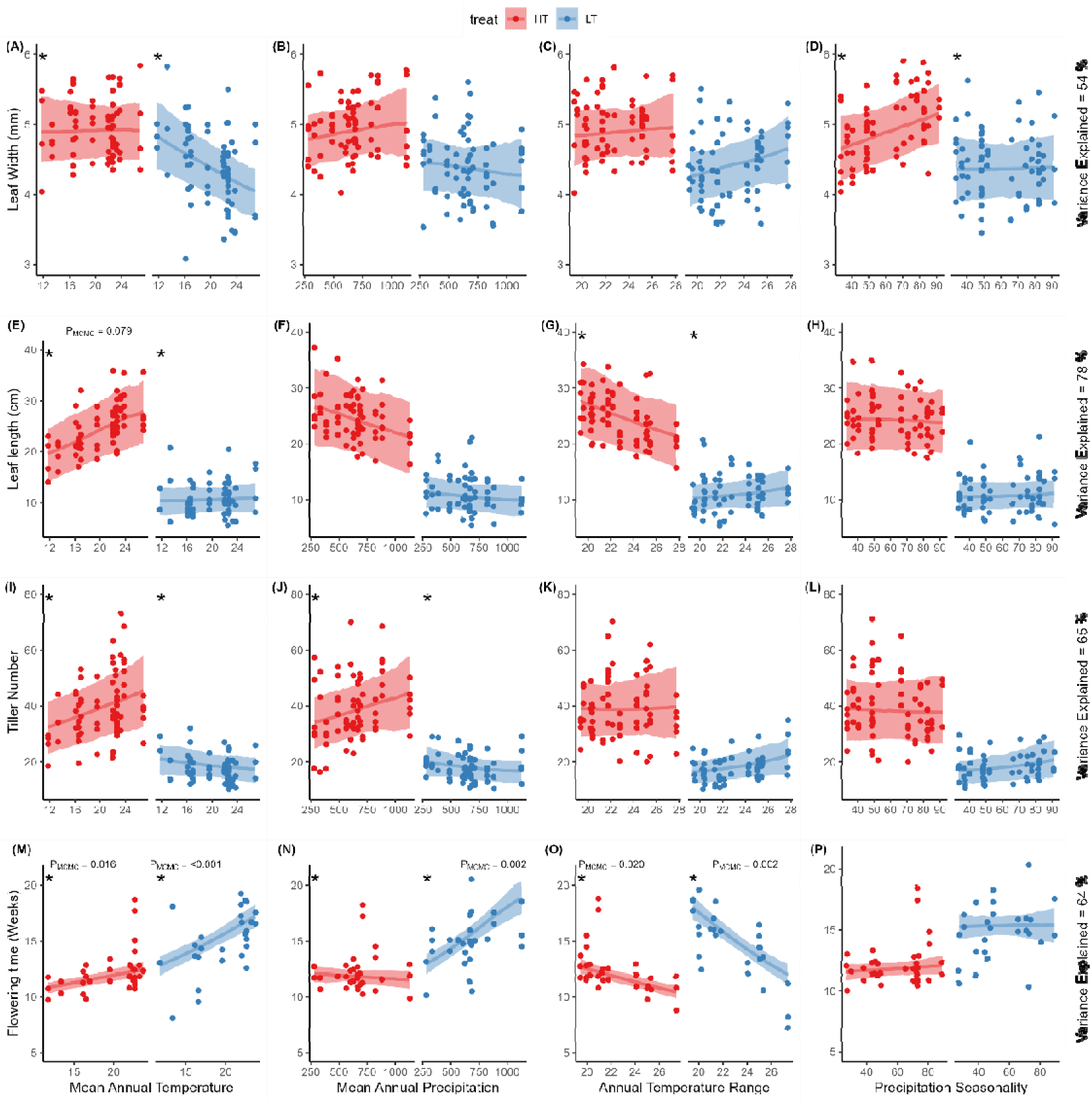
Conditional effects of site climate-at-origin on leaf morphology and reproduction. Traits: tiller number, leaf length, leaf width and flowering time. The solid lines and the shaded regions denote the mean and one standard error, respectively, of the posterior predictive distribution of the fitted models. The two-sided posterior probabilities for each slope (p_MCMC_) and for each slope difference (p_Δ,MCMC_) have been indicated if they were less than 0.1. Numerical p values are displayed for slopes associated with each growth temperature and an (*) for slope differences. For all features, red and blue denote the hot and cold growth temperatures (30°C and 20°C), respectively. For an explanation of conditional effects, refer to the caption in Figure 3. Detailed information on the posterior distributions of model parameters is provided in Figure S3.

In sum, several traits showed relationships to MAT but, in most cases, this was only the case for plants grown at the lower growth temperature. By contrast, mean annual precipitation at the site of origin showed no significant relationship with any trait other than to flowering time.

### Additional effects of climate seasonality

Annual temperature range (ATR) and precipitation seasonality (PS), treated statistically as secondary effects via residualisation, were similarly related to just a handful of traits. We observed a positive relationship between precipitation seasonality and A_sat_ (Fig. 3D), as predicted (hypothesis 6), but no effect of ATR on A_sat_. We observed a negative relationship between precipitation seasonality and R_mass_, at least for LT-grown plants (Fig. 4P). This suggests that higher-than-expected precipitation variability (for a given MAT and MAP) enhances photosynthetic activity and reduces R_mass_ or, alternatively, it is in some sense advantageous to have higher A_sat_ and lower R_mass_ under these conditions. WUE was lower in plants from higher-ATR sites but only for plants grown at the lower growth temperature (Fig. 3K); note, no relationship with ATR was observed for either A_sat_ or g_s_. LMA was lower in plants from higher-ATR sites but only for plants grown at the lower growth temperature (Fig. 4G). Flowering time was negatively related to annual temperature range, regardless of growth temperature (Fig. 5O).

## Discussion

We grew fifteen accessions of *T. triandra* from widely contrasting biomes in a glasshouse experiment to test whether physiological and morphological traits reflected the temperature and rainfall regimes from where seeds were collected – referred to here as their climate-of-origin (COO). Two thermal regimes were imposed to approximate the temperature extremes at the coolest and warmest sites. We used Bayesian linear mixed models to perform multiple regression with treatment interactions, taking into account the sampling structure and non-constant variability of the data. This enabled estimates of the relationships between each trait and climate variable, while simultaneously controlling for effects of the other climate variables and each glasshouse temperature regime. In general, physiological and morphological traits only revealed weak signals of the temperature or rainfall regimes from where the seeds originated, whereas flowering time revealed strong relationships with COO.

The lack of clear climate-related signals for many traits, put together with (i) clear treatment effects and (ii) divergent trait × COO relationships (i.e. different relationship slopes) in several cases, suggests that the morphological and physiological traits in *T. triandra* respond strongly to prevailing growing conditions – they are highly plastic. Importantly though, trait responses were occasionally specific to one of the two fixed-temperature regimes, reflected in a growing-temperature × COO interaction. In short, traits during *vegetative* growth were highly plastic, with all fifteen *T. triandra* accessions seemingly acclimating to the immediate thermal regimes in which they were growing and expressing few of the phenotypic traits that are typically associated with plants adapted to their native ranges (e.g. higher LMA and N_area_ in plants from drier and cooler sites (Wright and Westoby, 2002; Wright, Reich and Westoby, 2003; Wright *et al*., 2005; Prentice *et al*., 2014; Dong *et al*., 2017; Wang *et al*., 2023) and narrower leaves in plants from drier and hotter sites (Gates, 1965; Leigh *et al*., 2017; Wright *et al*., 2017; Baird *et al*., 2021). The relationship between flowering time and COO was a notable departure from the physiological and morphological traits discussed above. Strong correlations evident under cool growing conditions suggested that phenology was more highly “programmed” than vegetative traits. This suggests that the switch from vegetative to reproductive development is an adaptive shift that has become genetically fixed among accessions from contrasting thermal and precipitation regimes. This finding supports previous evidence that phenological events such as the time of floral initiation are critical adaptation mechanisms for survival in natural ecosystems (Franks *et al*., 2007; Lesica & Kittelson, 2010; Fournier-Level *et al*., 2022).

We showed that mean annual temperature was a stronger driver of trait variation in *T*. *triandra* than mean annual precipitation. We speculate that this is because temperature is an inescapable abiotic factor whereas grasses have multiple strategies that allow them to survive periods of low precipitation, including foliar senescence to allow meristems and roots to remain alive, and periods of dormancy interspersed with opportunistic growth (Turner, 1986; Volaire & Norton, 2006; Balachowski *et al*., 2016). This ability to survive dry periods by "shutting down" arguably reduces the selective pressure that precipitation may exert on functional traits as these grasses may not be actively growing or photosynthesising during these periods.

### Photosynthetic traits

We posited that plants adapted to colder sites would exhibit higher rates of photosynthesis and respiration compared with those from warmer sites, particularly when grown near their optimal temperatures of 20°C (Hypothesis 1). This idea was only weakly supported, with plants grown in cool conditions photosynthesising slightly faster when they were selected from cool-climate zones (Fig. 3A). On the other hand, rates of photosynthesis and respiration were highly responsive to the imposed temperature treatments, being generally faster for all accessions in warm than in cool glasshouses (Fig. 2). Similarly, at 20°C, warm-site plants had lower PNUE than plants from cool sites, reinforcing the hypothesis that C_4_ metabolism operates more efficiently at sub-optimal temperatures in accessions adapted to cool-temperate conditions. This generally indicates a greater thermal plasticity in cool-climate *T*. *triandra* accessions, which might go part way to explaining the expansion of *T. triandra* into the cool-temperate southern and elevated regions where mean annual temperatures are only ∼12°C. Supporting this observation, a range of plants from warm environments typically have higher thermal optima for photosynthesis (Berry & Bjorkman, 1980; Hikosaka *et al*., 2006) and thermal acclimation of photosynthesis is less effective in warm-adapted species (Way & Yamori, 2014). Accessions of *T. triandra* originating from warmer climates would therefore be predicted to perform poorly at suboptimal temperature conditions. Our findings do not reveal similar trends in relation to rainfall or seasonal variability, consistent with reports that plants from colder origins with shorter growing seasons generally exhibit higher photosynthetic rates when grown under common conditions (Benowicz, Guy and El-Kassaby, 2000; Kelly, Fletcher and Barney, 2020).

### Leaf dark respiration

We found no significant relationship between respiration and COO but, unsurprisingly, observed a significant overall reduction in respiration rates for all plants grown at the low growth temperature. Respiration is closely related to energy demand for biosynthesis, cellular maintenance and turnover of macromolecules required for growth and is highly temperature dependant (Atkin & Tjoelker, 2003; Atkin *et al*., 2015). Therefore, it acclimates with high sensitivity to prevailing environmental conditions, particularly temperature (Atkin & Tjoelker, 2003), and might be considered to be less intrinsically linked to the climatic origin of accessions within an individual species such as in this study. In general, acclimation of both photosynthesis and respiration seemingly masked the ‘memory’ of any adaptive phenomena that characterised each accession at its COO.

### Leaf economic traits

N_area_ showed no relationship with site precipitation (Hypothesis 2). This was unexpected because previous regional and global analyses of C_3_ species have shown a general tendency for species at drier sites to have higher N_area_, as predicted under “least-cost theory” (Wright *et al*., 2003; Prentice *et al*., 2014; Dong *et al*., 2020, 2022) where higher N_area_, connoting higher carboxylation capacity, allows species to operate at lower leaf-internal CO_2_ concentration during photosynthesis (c_i_), and to fix more C per unit water lost in transpiration. More simply, higher N_area_ has potential to drive higher WUE. We note that higher N_area_ at lower MAP can also be observed in the grasses-only component of the global “Glopnet” dataset (Wright et al 2004). So, why was this tendency not observed here? We offer two prospective explanations. First, if carboxylation capacity (and hence leaf N_area_) is rather easily “tuned” to prevailing growth conditions (Dong et al 2020), plants growing under common, well-watered conditions (as here) would show no systematic N_area_ variation in relation to precipitation at COO, even if field-sampled plants were to show the expected trend. Second, it is possible that the prediction is wrong, in that it applies to C_3_ species but not grass species with C_4_ photosynthesis, for which c_i_ lacks clear meaning due to their ability to concentrate CO_2_ around the site of carboxylation in the bundle sheath (Von Caemmerer & Farquhar, 1981).

Our other hypothesis for N_area_ was also not supported. We expected higher N_area_ at colder sites (Hypothesis 2), on the basis that more Rubisco is required at lower temperatures to generate a given carboxylation capacity (Prentice et al 2014) allowing for continued metabolic activity and growth under low temperatures (Reich & Oleksyn, 2004). There is clear evidence for this being true for woody species (Prentice et al 2014; Dong et al 2017), and re-visiting the grass (Poaceae) data from “Glopnet” (Wright et al 2004) provides additional support: N_area_ and MAT were strongly and negatively correlated if considered on their own (Fig. S4A; r^2^ = 0.14, P = 0.002, n = 91 species) and remained significant once MAP was simultaneosly considered (Fig. S4B; P = 0.026). Again, the discrepancy between predicted and observed trends might relate to common growing conditions or perhaps to differences between C_3_ and C_4_ species. Field sampling and analysis of more C_4_ species would help resolve these issues.

The observation that LMA was not related to precipitation did not accord with our expectation (Hypothesis 3) which was based on the presumed biomechanical advantages of thick, tough leaves in dry, physically demanding situations (Wright & Westoby 2002). Further, the observation that LMA increased with increasing MAT (at least at the low growth temperature) was opposite to our prediction (Hypothesis 3). That prediction was based on cost-benefit theory (Wang *et al*., 2023) and previous empirical work for evergreen woody trees. That said, in another literature strand a case has been made for an advantage of higher LMA at hotter sites: in hot, high light situations, the higher thermal mass of higher LMA leaves may provide brief but important protection from overheating when wind speeds drop (Leigh *et al*., 2012). In the grass component of the Glopnet dataset, LMA and MAT were strongly negatively correlated when considered on their own (Fig. S4C; r^2^ = 0.14, P < 0.001) and only marginally negatively correlated once MAP was simultaneously considered (Fig. S4C; P = 0.051, Wright *et al*., 2004).

### Leaf width and length

Although the disadvantages of wide leaves in hot environments are well understood via leaf energy balance theory (Gates, 1968; Parkhurst & Loucks, 1972; Givnish & Vermeij, 1976; Okajima *et al*., 2012; Wright *et al*., 2017) and, as such, we predicted narrow leaves to be characteristic of plants from high-MAT sites, we observed no such relationship in our study. Nor did we observe any relationship between leaf width and MAP. Again, we ascribe the absence of a trait-climate relationship to the high degree of trait acclimation. That is, it seems likely that when *T. triandra* is grown under common conditions, variation is masked in traits that – under fields conditions – adapt local populations to high and very low temperatures, water limitation, high irradiation, and so forth. Another possible reason for the lack of patterning between leaf width and MAT here is that narrow leaves are also often found in cold environments (Baird *et al*., 2021) as they are more closely coupled with air temperature, reducing the risk of night-time freezing (Jordan & Smith, 1995; Wright *et al*., 2017) and allowing plants to achieve a higher photosynthetic rate and water-use efficiency under warm conditions and compensate for shorter growing periods (Gates, 1968; Okajima *et al*., 2012; Körner, 2016). Our findings suggest that leaf width may be highly plastic, capable of responding to immediate growing conditions rather than being strictly determined by genetic adaptation to climate. If these plants were measured under field conditions, where local conditions such as temperature, humidity, and water availability vary significantly, we may expect to see stronger traitlclimate relationships, as is observed in other species (Okajima *et al*., 2012; Wright *et al*., 2017; Gallaher *et al*., 2019; Baird *et al*., 2021).

In contrast to what was observed for leaf width, leaf length and MAT showed a positive relationship, at least at the higher growth temperatures (30°C). This is consistent with plants from warm regions outperforming the cool-climate accessions when grown close to their thermal optimum (30°C). Determining growth rates in a classical (destructive) growth analysis at 20°C and 30°C was not possible because of high levels of seed dormancy limiting the availability of sufficient replicates, but leaf lengths are consistent with *T. triandra* having expanded its range southwards since its arrival in Australia (Dunning *et al*., 2017).

### Flowering time

As predicted (Hypothesis 5) plants from cooler environments initiated flowering earlier than plants from warmer environments. This was true for plants at either growth temperature although the effect was sharpest at 20°C (LT). Note also that plants grown at 20°C generally took longer to reach flowering than those for which 30°C. These trends most likely represent “heat sum” (growing-degree day) effects, i.e. plants from colder environments have a lower heat sum for flowering initiation (Bernier *et al*., 1993). Genotypic differences in flowering time may also be driven by photoperiod sensitivity, as observed in *Panicum virgatum* (Quinby, 1972; Sanderson *et al*., 1999; Casler *et al*., 2011). For instance, moving genotypes from higher latitudes (with long summer daylengths) to lower latitudes (with shorter summer day lengths) may induce flowering (van Esbroeck *et al*., 2003). In our study, we maintained a 14-h photoperiod, which potentially triggered earlier flowering in plants originating from the colder, more southern regions where summer daylengths typically exceed 15 hours. Previous studies have shown that flowering time is generally linked to genetic adaptation to local climatic conditions, particularly in response to temperature and photoperiod of their home environments (Körner & Basler, 2010; Wilczek et al., 2010). We also observed that plants originating from drier environments flowered sooner, when grown at 20°C, and similar trends have been observed in numerous other species (Lesica & Kittelson, 2010; Kigel et al., 2011). The early flowering in plants from drier environments is likely an adaptive strategy to avoid drought stress while plants from regions with higher and more reliable precipitation delay flowering, allowing for longer periods of vegetative growth and greater seed set. Overall, the shorter flowering times observed in plants from cold and dry environments are likely to result from the necessity of these plants to complete their reproductive cycle within a narrow window of favourable environmental conditions (Chmura *et al*., 2019).

### Seasonality effects on plant traits

Plants from environments with higher-than-expected temperature variability (i.e., for a given MAT and MAP) exhibited traits that seemingly enhance their ability to respond rapidly to resource availability, such as lower LMA and WUE and shorter time to flowering. These findings support Hypothesis 6 in that these traits are generallty indicative of a faster leaf economic strategy, allowing plants to capitalise on periods when temperatures are suitable for growth (Reich et al., 2003; Poorter et al., 2009). For example, in cool temperate environments such as Tasmania, where our lowest-temperature accession originated and where there is a much shorter growing season, plants must respond quickly when temperatures are optimal. Plants from environments with greater than expected precipitation seasonality, for a given MAT and MAP, had higher rates of photosynthesis, which may be an adaptation to increase responsiveness to rainfall and cope with periods of inconsistent rainfall (Huxman et al., 2004).

### Conclusions

Our detailed investigation under controlled glasshouse conditions provides new insights into trait-climate relationships in *T. triandra* and extends far beyond the current literature, which has primarily focussed on population genetics (Hayman, 1960; Evans & Knox, 1969; Dunning et al., 2017, 2022; Ahrens et al., 2020). In contrast to expectations, we observed limited and inconsistent relationships between most functional/morphological traits and climate-of-origin (COO). While quantitative differences between accessions for the traits we report here might have been apparent if the fifteen accessions had been sampled in their natural environment, the significance of this study is that they were largely absent in plants grown under common conditions. Overwhelmingly, vegetative traits were highly labile and conformed to current growing conditions. This plasticity is seemingly a testament to the adaptability of *T. triandra* and its capacity to optimise trait values to prevailing environmental conditions. The observed clear link between flowering time and COO underscores the importance of phenology as a strongly encoded (‘fixed’) trait in this species. We propose that adaptation to climate in *T. triandra* is driven by a combination of high trait plasticity and the ability to synchronise reproduction with favourable conditions, particularly periods of high temperature, which is critical for growth in C4 grasses. Sampling the traits of this widespread species in natural situations across its range and comparing the data to those collected here would likely provide a deeper understanding of climate adaptation in this species.

## Acknowledgements

We thank Dr Alan Humphries from the Australian Pastures Genebank for advice on germinating *T. triandra* and for providing seed sources that helped make this project possible. IJW, VJ, LY, ES and TB acknowledge support from the Australian Research Council (CE200100015).

## Author contributions

VJ and BJA led project design, experimental work and manuscript preparation, with strong support from IJW. TB and ES assisted with data collection. LY and VJ led data analysis. All authors discussed results, contributed ideas and text to multiple versions of the manuscript, and approved the final version.

## Data availability

The full dataset will be made available in Supplementary Material upon acceptance

## Supplementary Tables and Figures

**Figure S1:**
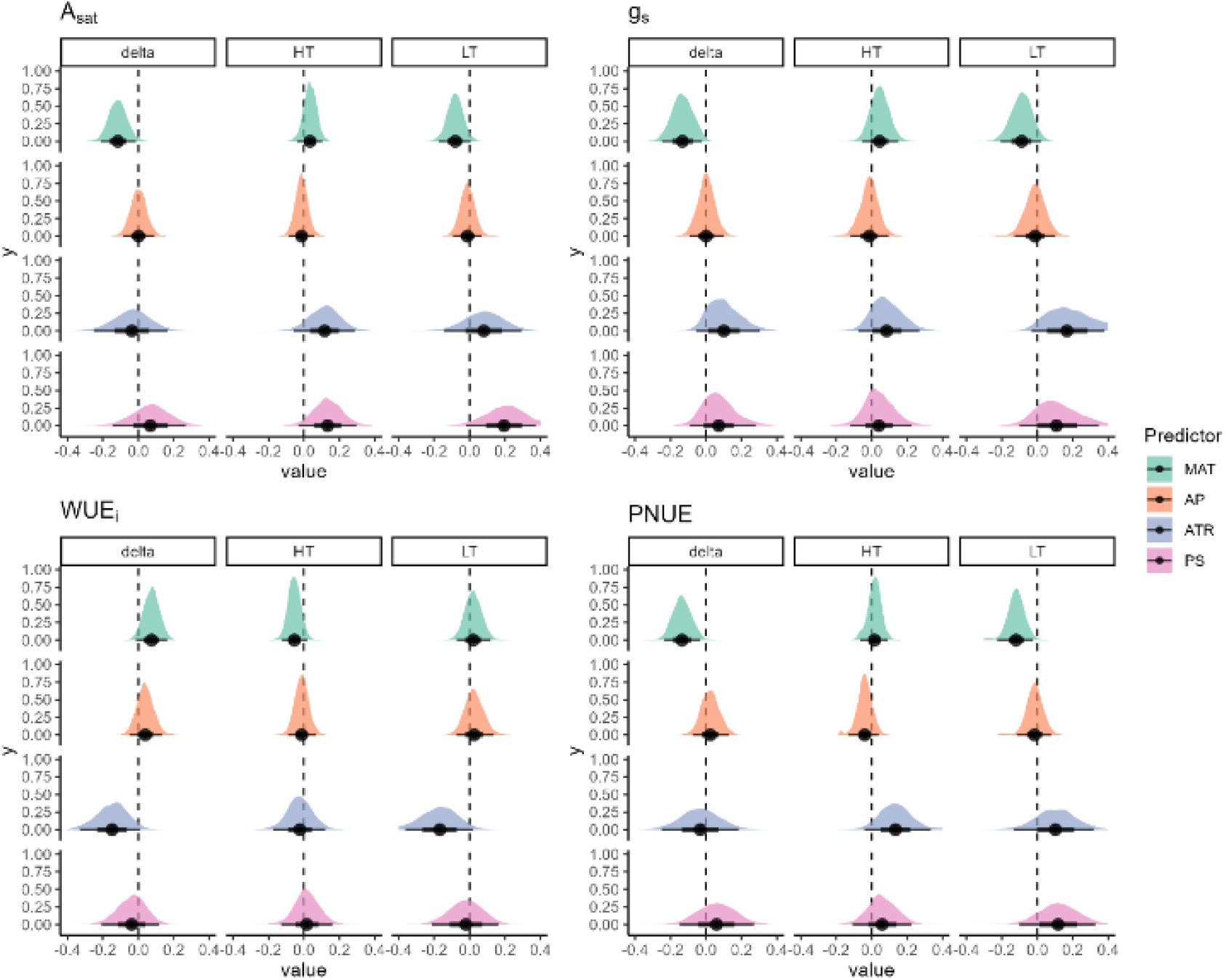
Effects of four climate predictors on plant physiological traits across growth temperatures. Posterior distributions for the linear effect of Mean Annual Temperature (MAT, green), Annual Precipitation (AP, orange), Annual Temperature Range (ATR, blue), and Precipitation Seasonality (PS, pink) on light saturated photosynthetic rate rate(A_sat_), stomatal conductance to water vapour (g_s_), intrinsic water use efficiency (WUE_i_, A_sat_:g_s_) and photosynthetic nitrogen use efficiency (PNUE, A_sat_:N_area_). Each panel displays effect estimates for high temperature (HT, 30°C) and low temperature (LT, 20°C) growing conditions, as well as the effect difference between the conditions (Δ). The coloured areas represent the posterior densities, with black dots and lines denoting the mean and 95% credible intervals, respectively.

**Figure S2:**
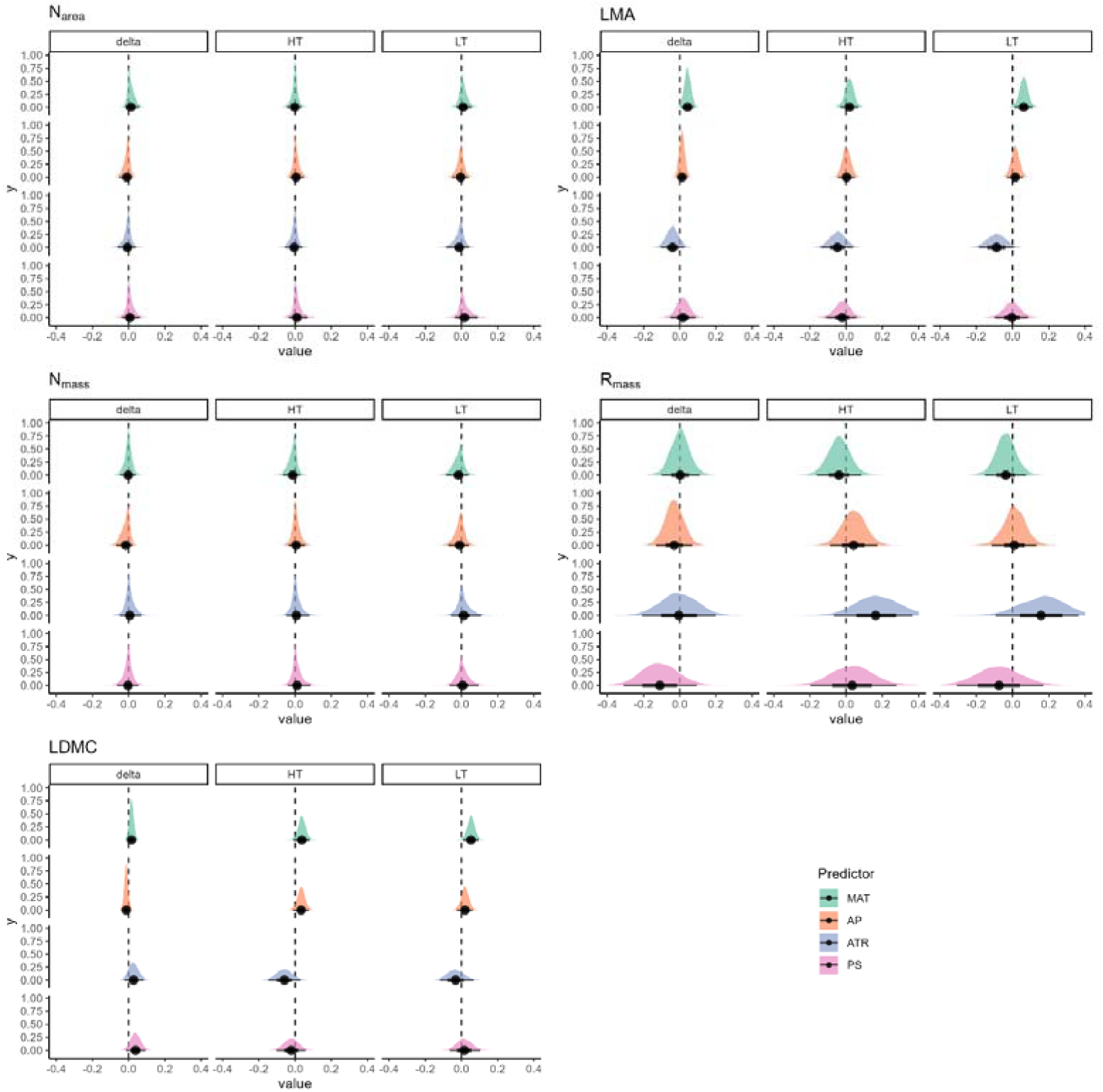
Effects of four climate predictors on leaf economic traits across growth temperatures. Posterior distributions for the linear effect of Mean Annual Temperature (MAT, green), Annual Precipitation (AP, orange), Annual Temperature Range (ATR, blue), and Precipitation Seasonality (PS, pink) on leaf Nitrogen per area (N_area_), leaf mass per area (LMA), leaf Nitrogen per mass (N_mass_), leaf dark respiration per mass (R_mass_) and leaf dry matter content (LDMC). Each panel displays effect estimates for high temperature (HT, 30°C) and low temperature (LT, 20°C) growing conditions, as well as the effect difference between the conditions (Δ). The coloured areas represent the posterior densities, with black dots and lines denoting the mean and 95% credible intervals, respectively.

**Figure S3:**
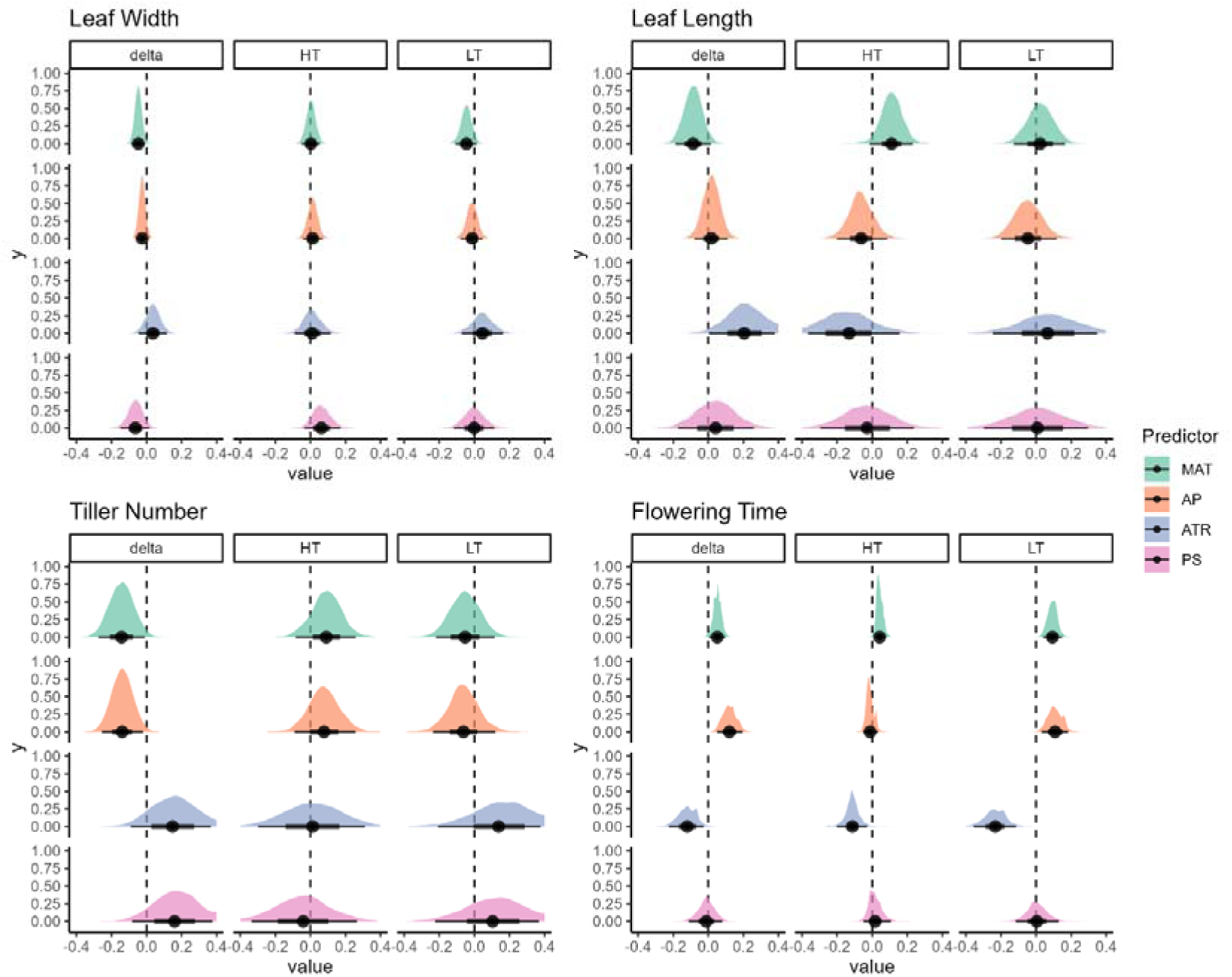
Effects of four climate predictors on leaf economic traits across growth temperatures. Posterior distributions for the linear effect of Mean Annual Temperature (MAT, green), Annual Precipitation (AP, orange), Annual Temperature Range (ATR, blue), and Precipitation Seasonality (PS, pink) on leaf width, leaf length, tiller number and flowering time. Each panel displays effect estimates for high temperature (HT, 30°C) and low temperature (LT, 20°C) growing conditions, as well as the effect difference between the conditions (Δ). The coloured areas represent the posterior densities, with black dots and lines denoting the mean and 95% credible intervals, respectively.

**Figure S4.**
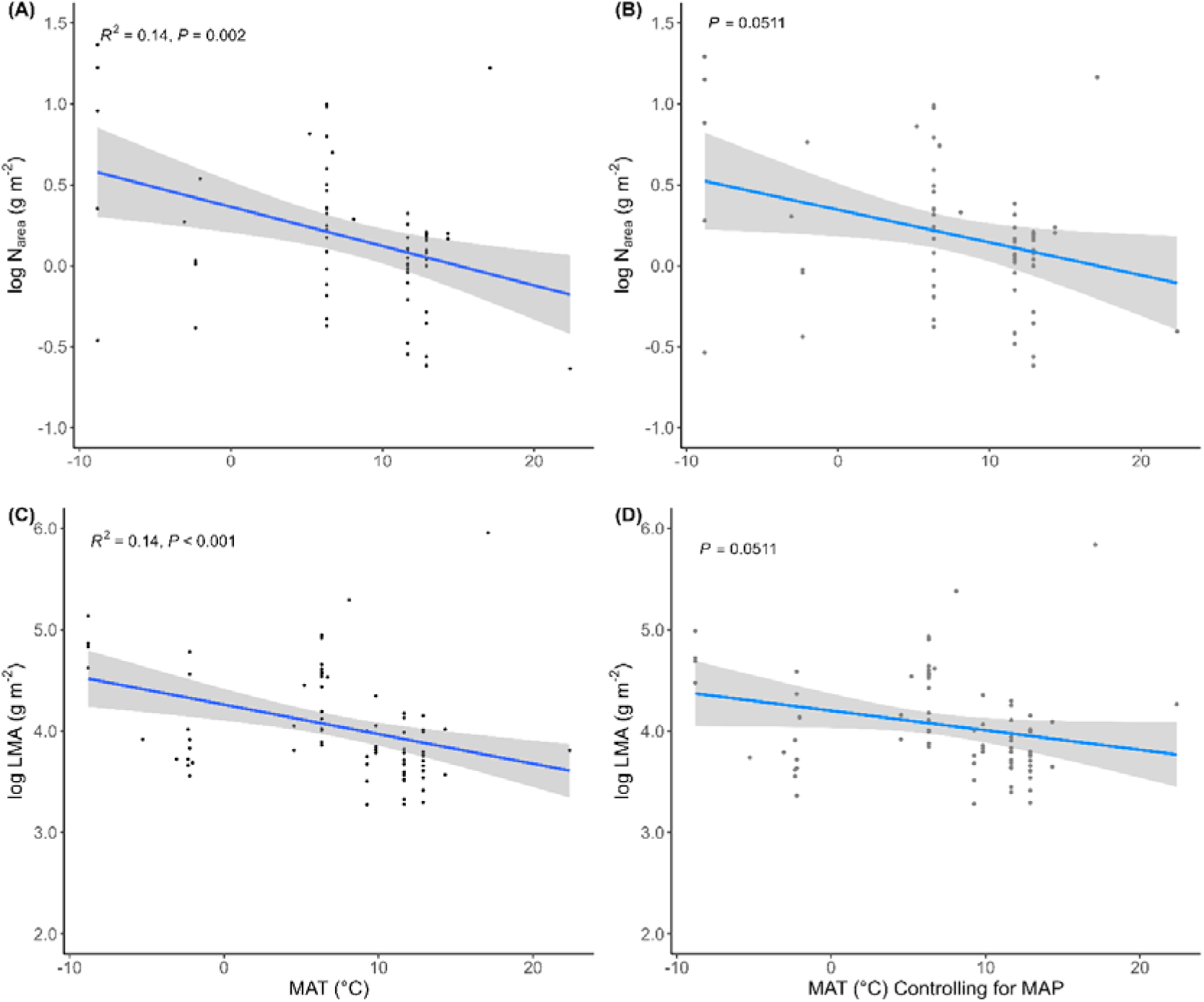
Trait – climate relationships. Relationship between mean annual temperature (MAT, °C) and (A) Leaf nitrogen per area (N_area_) and (C) leaf mass per area (LMA). (B) Partial residual plot for leaf nitrogen per area (N_area_) and (D) leaf mass per area (LMA) as a function of MAT while controlling for mean annual precipitation (MAP, mm). Blue regression lines represent model-predicted trends, while grey shaded areas denote 95% confidence intervals.

**Table S1:** Geographic origin and historical climate data for the fifteen Themeda triandra accessions included in this study.

| Accession | State | latitude | longitude | MAT | MAP | ATR | PS |
| --- | --- | --- | --- | --- | --- | --- | --- |
| Fumar | QLD | -17.13545 | 145.1834 | 22.7 | 823 | 18.1 | 105.1 |
| Forbes | NSW | -33.4076 | 147.9654 | 17 | 490 | 27 | 10.2 |
| Rainbow Valley | NT | -24.2376 | 133.5536 | 22.1 | 281 | 31.2 | 50.2 |
| Palm Valley | NT | -23.9666 | 132.75 | 22 | 333 | 30.1 | 54.9 |
| Virginia | SA | -34.8538 | 138.6205 | 16.2 | 668 | 18.7 | 51.2 |
| Hotspur | VIC | -37.9171 | 141.6471 | 13.4 | 686 | 18.7 | 36.6 |
| Mt Coolon | QLD | -21.4055 | 147.3506 | 23.2 | 605 | 23.2 | 69.1 |
| Dalby | QLD | -27.1148 | 151.1817 | 19.6 | 627 | 24.5 | 43.7 |
| Conjuboy | QLD | -18.9926 | 144.6917 | 22.5 | 683 | 21.5 | 108.4 |
| Woodstock | QLD | -19.6155 | 146.9620 | 23.8 | 883 | 18.7 | 98.6 |
| Nile | TAS | -41.6461 | 147.3287 | 11.8 | 564 | 19.4 | 24.4 |
| Mount Fox | QLD | -19.0013 | 145.4731 | 22.7 | 712 | 20.7 | 94.1 |
| Pannawonica | WA | -21.6449 | 116.3233 | 26.9 | 387 | 27.6 | 108.6 |
| Sydney | NSW | -33.6881 | 151.0624 | 16.6 | 1129 | 20 | 23.8 |
| Marlborough | QLD | -22.8777 | 149.7922 | 22.7 | 666 | 20.4 | 58.2 |

Climate data (1981–2010) are from the CHELSA (Climatologies at high resolution for the earth's land surface areas) data of downscaled model output temperature and precipitation estimates of the ERA-Interim climatic reanalysis to a high resolution of 30'arcsec. MAT, Mean Annual Temperature (°C); MAP, Mean Annual Precipitation (mm); ATR, Annual Temperature Range (°C); PS, Precipitation Seasonality (Coefficient of Variation).

